# Auditory Target Detection Enhances Visual Processing and Hippocampal Functional Connectivity

**DOI:** 10.1101/2020.09.19.304881

**Authors:** Roy Moyal, Hamid B. Turker, Wen-Ming Luh, Khena M. Swallow

## Abstract

Though dividing one’s attention between two input streams typically impairs performance, detecting a behaviorally relevant stimulus can sometimes enhance the encoding of unrelated information presented at the same time. Previous research has shown that selection of this kind boosts visual cortical activity and memory for concurrent items. An important unanswered question is whether such effects are reflected in processing quality and functional connectivity in visual regions and in the hippocampus. In this fMRI study, participants were asked to memorize a stream of naturalistic images and press a button only when they heard a predefined target tone (400 or 1200 Hz, counterbalanced). Images could be presented with a target tone, with a distractor tone, or without a tone. Auditory target detection increased activity throughout the ventral visual cortex but lowered it in the hippocampus. Enhancements in functional connectivity between the ventral visual cortex and the hippocampus were also observed following auditory targets. Multi-voxel pattern classification of image category was more accurate on target tone trials than on distractor and no tone trials in the fusiform gyrus and parahippocampal gyrus. This effect was stronger in visual cortical clusters whose activity was more correlated with the hippocampus on target tone than on distractor tone trials. In agreement with accounts suggesting that subcortical noradrenergic influences play a role in the attentional boost effect, auditory target detection also caused an increase in locus coeruleus activity and phasic pupil responses. These findings outline a network of cortical and subcortical regions that are involved in the selection and processing of information presented at behaviorally relevant moments.

## 1. Introduction

Attention can be allocated not only to spatial locations or stimulus features (Carrasco, 2011), but also to information presented at particular points in time (Rohenkohl et al., 2011; Nobre & van Ede, 2018). A growing literature shows that perceptual processing is enhanced when events change in meaningful ways (e.g., Jefferies & Di Lollo, 2019) and when they require a response (Swallow & Jiang, 2010; Makovski et al., 2013; Yebra et al., 2019; Clewett et al., 2020). Detecting a behaviorally relevant item, for instance, can improve memory for concurrently presented but otherwise unrelated information (the attentional boost effect, or ABE; Swallow & Jiang, 2010), even when it is task-irrelevant (Broitman & Swallow, 2020; Swallow & Jiang, 2014; Turker & Swallow, 2019). Target detection can also increase visual adaptation, lexical priming, and affective evaluation of concurrently presented items (Pascucci & Turatto, 2013; Spataro et al., 2013; Schonberg et al., 2014; Swallow & Atir, 2018). The beneficial effects of presenting information at the same time as a target can be contrasted with those commonly observed in attention tasks that require participants to select among different sources of information. Under these conditions, competitive interactions within and between regions are associated with reduced processing of unselected information (e.g., in visual cortex when monitoring auditory rather than visual stimuli, Johnson & Zatorre, 2005; of multi-voxel patterns associated with task irrelevant categories when searching for a pre-specified category of objects in natural images, Seidl et al., 2012; in parts of topographically organized visual cortex in uncued regions of a visual display during a search task, Silver et al., 2007). However, despite the extensive evidence that the selection of one item (such as an auditory tone) reliably boosts behavioral indices of background item (such as a visual scene) processing, little is known about its neural basis. Guided by previous empirical work, this project used fMRI to study the neurophysiological basis of the effects of target detection on visual processing and memory in the ABE paradigm. Specifically, we examined whether it enhances the quality of representations in—and communication between—regions involved in episodic encoding.

Consistent with prior work demonstrating that norepinephrine (NE) increases neural gain in response to behaviorally relevant events or task boundaries (Aston-Jones & Cohen, 2005; Bouret & Sara, 2005; Lee et al., 2018), the ABE could reflect the phasic firing of the locus coeruleus (LC) in response to behaviorally relevant events (Swallow & Jiang, 2013). The LC, a brainstem structure whose activity briefly increases in response to changes in a task or in the environment (Clewett et al., 2020; Sara, 2009) and facilitates episodic encoding (Takeuchi et al., 2016), is the main source of NE in the brain. Phasic LC responses correlate with pupil diameter (Murphy et al., 2014; Joshi et al., 2016) and are associated with target detection and orienting (Aston-Jones et al., 1994; Breton-Provencher & Sur, 2019). Though previous studies suggest a relationship between LC activity and the ABE (Swallow et al., 2019; Yebra et al., 2019), they utilized indirect measures (pupil size) or a probabilistic atlas to identify the LC in participants, making it difficult to pinpoint the source of the modulatory signals (cf. Wang & Munoz, 2015). This is of particular concern because the small size of the LC and its location near the fourth ventricle, a source of physiological noise in fMRI, increase the potential for mislocalization and for the inclusion of spurious signals in estimates of LC activity (Turker et al., 2021). We therefore utilized structural MRI T1 sequences that increase contrast for the high concentrations of neuromelanin in the LC (Keren et al., 2009) to improve our ability to localize the LC in individual participants relative to a probabilistic atlas (Turker et al., 2021). We also employed multi-echo EPI with multi-echo independent components analysis (ME-ICA) and TE-dependent BOLD signal classification (Kundu et al., 2013) to reduce the contributions of noise sources to our data.

The effects of target detection on episodic encoding, visual adaptation, and lexical priming (e.g., Broitman & Swallow, 2020; Pascucci & Turatto, 2013; Spataro et al., 2013; Turker & Swallow, 2019) suggest that it should improve the quality of representations in perceptual and episodic encoding regions. Such effects would also be expected if target detection increases neural gain, enhancing the signal to noise ratio of activity in impacted regions (e.g., Aston-Jones & Cohen, 2005). However, fMRI investigations of the ABE have provided little insight into the mechanisms by which it modulates neural processing. Prior work has shown that target detection broadly increases the BOLD signal in regions not directly involved in processing the target stimulus (e.g., auditory target detection boosts activity in V1; Jack et al., 2006; Swallow et al., 2012). These studies did not include baseline trials, however, making it unclear whether the reported effects reflect target-related facilitation or distractor-related inhibition. Moreover, differences in the magnitude of the hemodynamic response do not, on their own, reflect differences in processing quality (cf. Albers et al., 2018). This study therefore incorporated baseline trials and used multivoxel pattern analysis to test whether target detection enhances the quality of processing (Mahmoudi et al., 2012). Attention-related enhancements in processing quality often coincide with changes in the amount or spread of decodable representational information in perceptual regions (e.g., Zhang et al., 2011); in the medial temporal lobe, increases in BOLD magnitude and decoding accuracy (e.g., Chadwick et al., 2010; Chadwick et al., 2011) have been linked to episodic encoding and recall. Nonetheless, the relationship between auditory target detection and the quality of visual processing and encoding remains underexplored.

Prior research also leaves the possibility that target detection in the ABE paradigm affects coordination among different brain regions unexplored. Better working memory (Gazzaley et al., 2004) and episodic encoding (Ranganath et al., 2005) are associated with enhanced functional connectivity (Friston, 2011) between the hippocampus (HPC) and visual areas. Findings of enhanced short-term memory (Li et al., 2018; Makovski et al., 2011) and episodic encoding (Broitman & Swallow, 2020; Leclercq & Seitz, 2014; Mulligan et al., 2021; Turker & Swallow, 2019) with the ABE thus suggests that it should increase functional connectivity between these regions. Phasic LC activation also is temporally coordinated with the HPC, anterior cingulate cortex (ACC), and other prefrontal regions (cf. Sara, 2015) and may lead to the dynamic reconfiguration of cortical functional networks (Bouret & Sara, 2005; Shine et al., 2018; Li et al., 2019). Disruptions to existing population firing patterns created by phasic LC activation could also facilitate the formation of patterns that represent behaviorally relevant information (Moyal & Edelman, 2019). While these findings suggest that auditory target detection should trigger an increase in visuo-hippocampal connectivity, this possibility has yet to be examined directly. It may also be possible that the anterior and posterior hippocampus (aHPC and pHPC, respectively) are differently impacted by target detection. Relative to pHPC, aHPC is more strongly associated with episodic memory encoding (relative to spatial memory encoding), is associated with more generalized (less detailed) representations of events, shows stronger functional connectivity with fusiform gyrus (FG) and medial versus lateral aspects of entorhinal cortex, and may have greater concentrations of NE receptors (Frank et al., 2019; Persson et al., 2018; Poppenk et al., 2013; Gage & Thompson, 1980). However, non-human animal research also suggests that the LC may play a role in modulating episodic memory formation in pHPC (Kempadoo et al., 2016; Wagatsuma et al., 2018). We therefore investigated the effects of auditory target detection in the ABE paradigm on the functional connectivity of aHPC and pHPC to visual areas.

To summarize, we used multi-echo fMRI to characterize the neural correlates of target detection in the ABE, specifically examining responses of the visual cortex, HPC, and LC to images presented on their own or with auditory target or distractor tones (Swallow & Jiang, 2010; Swallow et al., 2012). In contrast to previous work, we expected target detection to increase (1) phasic pupil responses and activity in individually defined LC; (2) the ability to classify patterns of BOLD activity associated with different categories of images; and (3) functional connectivity between visual regions and HPC. We found evidence supporting each of these hypotheses.

## 2. Materials and Methods

### 2.1 Participants

21 right-handed individuals (15 female, 6 male, 19-40 years old, M = 21.48, SD = 4.86) participated in the study. They were screened for non-MRI compatible medical devices or body modifications, claustrophobia, movement disorders, pregnancy, mental illness, use of medication affecting cognition, and color blindness. Consent was obtained at the beginning of the session and participants were debriefed at the end. All procedures were approved by the Cornell University review board. Sample size was based on a previous study examining the effect of auditory target detection on visual cortical activity (Swallow et al., 2012), which reported effect sizes for a peak signal difference following target and distractor auditory tones of *Cohen’s f* > 1.037. A sample size of 20 was selected to ensure that smaller effects between conditions and in other measures of connectivity and classification could be detected. With a sample of 20 and false positive rate of .05, a traditional one-way (three levels) repeated measures analysis of variance has a power of .95 to detect an effect of *Cohen’s f* > .378 (calculated using G*Power; Faul et al., 2007).

Two participants responded to the wrong tones on some scans so that some images were paired with both target and distractor tones (one of these participants also did not complete the memory test). Because these participants were performing the target detection task (with the wrong tone) their data for these scans were recoded and included in analyses of detection task performance and in the univariate analyses (which had an N of 21). However, these two participants were excluded from analyses that depended on balancing the number of trials across conditions (image classification and functional connectivity, which fed into an image classification analysis), resulting in an N of 19 for these analyses. They were also excluded from analyses involving the memory test. One additional participant did not complete the memory test due to a fire alarm, leaving 18 participants for all analyses involving memory data.

### 2.2 MRI and Pupillometry Data Acquisition

Magnetic resonance imaging was performed with a 3T GE Discovery MR750 MRI scanner (GE Healthcare, Milwaukee, WI) and a 32-channel head coil at the Cornell Magnetic Resonance Imaging Facility in Ithaca, NY. Participants laid supine on the scanner bed with their head supported and immobilized. Ear plugs and headphones (MR confon GmbH, Germany) were used to reduce scanner noise, allow the participant to communicate with the experimenters, and present auditory stimuli during the tasks. Visual stimuli were presented with a 32” Nordic Neuro Lab liquid crystal display (1920×1080 pixels, 60 Hz, 6.5 ms g to g) located at the head of the scanner bore and viewed through a mirror attached to the head coil.

Anatomical data were acquired with a T1-weighted MPRAGE sequence (TR = 7.7 ms; TE = 3.42 ms; 7° flip angle; 1.0 mm isotropic voxels, 176 slices). A second anatomical scan utilized a neuromelanin sensitive T1-weighted partial volume turbo spin echo (TSE) sequence (TR = 700 ms; TE = 13 ms; 120° flip angle; 0.430 x 0.430 mm in-plane voxels, 10 interleaved 3.0 mm thick axial slices; adapted from Keren et al., 2009). Slices for the TSE volume were oriented perpendicular to the long axis of the brain stem to provide high resolution data in the axial plane, where dimensions of the LC are smallest, and positioned to cover the most anterior portion of the pons. Multi-echo echo planar imaging (EPI) sequences were used to acquire functional data during the four task runs (TR = 2500 ms; TEs = 12.3, 26.0, 40.0 ms; 83° flip angle; 3.0 mm isotropic voxels; 44 slices). In addition to the task runs, all participants also completed a single resting state scan with their eyes open and the lights on (612s; TR = 3.0 s; TEs=13, 30, 47 ms). Resting state data are reported elsewhere (Turker et al., 2021) but were used for this study (see 2.5.9 *LC Functional Connectivity*).

During the scans, pupil size and gaze location were acquired using an EyeLink 1000 Plus MRI Compatible eye tracker (SR-Research, Canada) for all but two participants (1000 Hz, right eye). After the participant was positioned in the scanner, mirrors were adjusted to bring the eye into view of the camera. Immediately prior to the resting state scan, thresholds defining pupil and corneal reflectance were automatically adjusted and a nine-point calibration routine was performed to determine the parameters needed to estimate gaze position. Calibration was validated and adjusted as necessary prior to each scan that included eye data measurement. On task runs, participants were instructed to fixate the central dot and minimize blinking.

### 2.3 MRI Data Preprocessing

All EPI data were denoised and processed using the standard ME-ICA pipeline, except as indicated (meica.py, Version 3.2, beta 1; Kundu et al., 2012; Kundu et al., 2013). First, the MPRAGE volume was skull stripped using FSL v5.0 BET (b = 0.25). After matching the obliquity of the anatomical volume and EPI time series, motion was estimated from the first echo time series using 3dvolreg and the third volume as the target. Third, all EPI data were despiked and slice time acquisition differences were corrected using 3dTshift. Fourth, for each echo time series, the first two volumes were dropped and the remaining EPI data were registered to the third volume. Baseline intensity volume (s0), the t2* map volume (t2*), and the optimal combination volume time series were then calculated. Fifth, registration and alignment transforms were applied to the EPI data and the pre-equilibrium volumes dropped in one step to align the data with the individual anatomical volume in its original acquisition space. Sixth, EPI data were denoised to identify and separate BOLD components from non-BOLD components (Kundu et al., 2013). BOLD components were recombined to create the denoised data sets that were used in subsequent analyses. Finally, denoised EPI data were spatially aligned to the MNI N27 atlas for volume-wise group level analyses.

### 2.4 Region of Interest Identification

Individual MPRAGE scans were submitted to FreeSurfer’s segmentation and surface-based reconstruction software (recon-all v5.3; surfer.nmr.mgh.harvard.edu; Dale et al., 1999; Fischl et al., 1999) to label voxels according to each individual’s anatomy. Labels for the fourth ventricle (4V), hippocampus (HPC), motor cortex (MC), planum temporale for auditory cortex (AC), primary visual cortex (V1), secondary visual cortex (V2), fusiform gyrus (FG), and parahippocampal gyrus (PG) were extracted and converted to volumetric ROIs using FreeSurfer and AFNI tools (Cox, 1996; Cox & Hyde, 1997; Gold et al., 1998). Separate ROIs were created for the left and right hemispheres. In addition, the HPC ROIs were divided into anterior and posterior portions at the anterior-posterior coordinate of their center of mass (aHPC, pHPC), to account for possible differences in their connectivity patterns and function (Fanselow & Dong, 2010; Poppenk et al., 2013), Detailed methods for identifying the LC are described in Turker et al. (2021). Briefly, individual MPRAGE scans (including skull) were aligned to the individual normalized T1-weighted neuromelanin scan. After extracting the brainstem from the nmT1, correcting image intensity, and setting the false color palette to the predefined range, candidate LC voxels could be visually distinguished from nearby regions (Figure 3A, S3). Bilateral LC ROIs were then hand-drawn by two tracers (voxel size: 0.43 x 0.43 x 3.0 mm). Voxels included as LC by both raters were kept, resampled to 3 mm isotropic voxels, and spatially aligned to the individual EPI and MPRAGE. This procedure resulted in an average of 4.5 voxels per participant (SD = 1.9). Finally, we also created an ROI for the ACC, based on its connections with the LC (e.g., Ennis et al., 1998). The association test map for the ACC was downloaded from Neurosynth (neurosynth.org) and thresholded (*z* = 6) to retain an ROI covering only putative ACC.

### 2.5 Experimental Design and Statistical Analysis

#### 2.5.1 Stimuli

288 full color images of faces (48 female, 48 male), objects (48 cars, 48 chairs), and outdoor scenes (48 beaches, 48 forests) were acquired from personal collections and publicly available online databases (vision.stanford.edu/projects/sceneclassification/resources.html; Huang et al., 2007; Huang & Learned-Miller, 2014; Xiang et al., 2014). 24 images from each subcategory were used in the encoding task (eight in each tone type condition), each presented once per run (four repetitions in total). The rest of the images were used as foils in the recognition test. Scrambled images were generated from these photographs by dividing them into 256 tiles and shuffling their locations. The mean and variance of pixel intensity (luminance) was matched across images using the SHINE toolbox (Willenbockel et al., 2010). The presentation of simple auditory stimuli with complex naturalistic images is standard for this paradigm (e.g., Swallow & Jiang, 2010; Swallow & Jiang 2012; Swallow et al., 2019; Swallow, Makovski & Jiang, 2012) and allows for the separation of the effects of selection on processing the target and distractor tones from the impact of auditory target detection on visual stimuli processing.

#### 2.5.2 Design and Procedure

In the four functional runs (407.5 s each) participants continuously performed simultaneous image encoding and target detection tasks. On each 1.25 s long trial, one image (7 x 7 visual degrees; 256 x 256 pixels) was presented for 625 ms and immediately followed by another image for another 625 ms. On most trials, both images were scrambled (no-task trials, 164/run). On task trials (144/run), the first image was a photograph and the second was a scrambled version of that photograph. Participants attempted to memorize the photograph for a later memory test. The inter-trial interval was 0 ms, ensuring that there was an image on the screen throughout the task and that it changed every 625 ms (Figure 1A). A red fixation dot (0.25 visual degree diameter) appeared at the center of the screen throughout the task, including 7.5 s of pre-task fixation and 15 s of post-task fixation at the beginning and end of each run, respectively.

**Figure 1.**
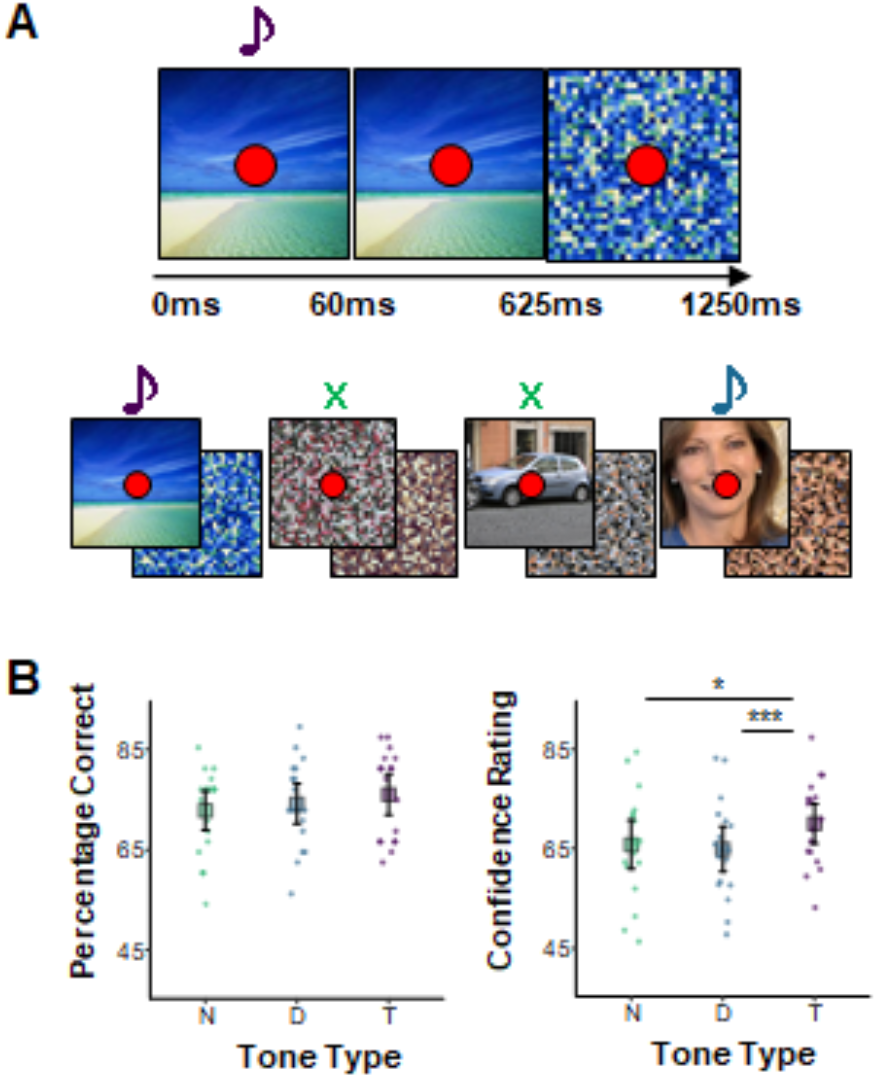
Experimental paradigm and the effect of tone type on subsequent memory and whole brain activity. **A:** Trial and trial series of the image encoding and target detection task performed during scanning. Each photograph was presented for 625ms and followed by one or more scrambled images. The photographs belonged to one of six categories and could be accompanied by a target tone (purple) that warranted a button press, a distractor tone (blue) that did not, or no tone (green). Tones were 60ms long. **B:** Left: percentage of correctly recognized images during a post-scanning two alternative forced choice recognition test. Right: confidence ratings (continuous scale, 0-100) for correctly remembered images. Each point represents a single participant’s mean. Large squares are centered on the sample mean and error bars indicate 95% confidence intervals around the mean. *p < .05, **p < .01, ***p < .001 in a Holm-corrected general linear test comparing conditions (see 2.6 in *Materials and Methods*).

On some task trials a high (1200 Hz) or low (400 Hz) pitched auditory tone (60 ms duration) was played over the headphones at the same time an image was presented (0 ms stimulus onset asynchrony). If the tone was the pre-specified target pitch participants pressed a button with their right thumb. Participants made no overt response if the tone was not the pre-specified target pitch (distractor) or if no tone was presented on that trial. The tone assigned to the target condition was counterbalanced within participants by switching it halfway through the encoding task (e.g., high, high, low, low). Participants were told which tone was the target tone at the beginning of each run. The starting target tone was counterbalanced across participants: Half started with the high tone as the target and half started with the low tone as the target. Tone condition (target, distractor, no tone) was held constant for each image. Thus, if a high tone was assigned to a given image on the first two runs, it was switched to a low tone on the latter two runs. An equal number of task trials was assigned to each tone type (target, distractor, and no tone), ensuring that results cannot be attributed to the salience or relative frequency of target occurrences. Tones were never presented on no-task trials. Sound levels were adjusted during the MPRAGE scan to ensure participants could hear both tones during scanning. Participants practiced the task with a different set of images before entering the scanner.

After scanning participants completed a two alternative forced choice recognition test on the images. On each trial two images were presented on the screen, one on the left side and one on the right. One of the images was presented during the encoding and detection task and the other was a new image. Participants selected the ‘old’ photograph by pressing one of two keys on the keyboard. Participants were then prompted to report their confidence by clicking on a line that appeared below the images. Participants were told to click on the far-left side of the line if they were guessing, the far-right side of the line if they were absolutely confident that they were correct, and at points in between to reflect degrees of intermediate levels of confidence. These were coded on a scale from 1 (lowest rating) to 100 (highest rating). A green + or a red - appeared next to indicate their accuracy. This procedure resulted in a 6 x 3 design, with *image type* (female face, male face, beach, forest, car, chair) and *tone type* (no tone, distractor tone, target tone) as within-participants factors. There were 32 trials per image-by-tone condition for a total of 576 task trials over four runs. Each run started with 7.5 s of fixation and ended with 15 seconds of fixation. Trial order and spacing were optimized using the AFNI function make_random_timing to produce four sequences that minimized the amount of unexplained variance in a simulated task. Task trials were separated by 0-12 non-task trials.

#### 2.5.3 Behavioral Data Analysis

To examine the effect of tone type on memory we fit a binomial generalized linear mixed effects model (Bates et al., 2015) to the recognition accuracy data with tone type as a fixed effect and with random intercepts for participant, old image, and new image. A linear mixed effects model was fit to participants’ recognition confidence ratings for correctly recognized images, with the same variables and random intercepts.

#### 2.5.4 Tonic and Phasic Pupil Size Estimation and Analysis

Before estimating tonic and phasic pupil dilation on a trial-by-trial basis, pupil data for each participant and task run were preprocessed using the EyeLink DataViewer application (SR-Research, Canada), the FIRDeconvolution toolbox (Knapen et al., 2016), and custom routines. In brief, following the procedure outlined in Knapen et al. (2016), linear interpolation was used to estimate pupil size during blinks flagged by the EyeLink software and extended to include 100 ms margins before and after the blink. High-pass (0.1 Hz) and low-pass (10 Hz) Butterworth filters were applied, after which the data were down-sampled from 1000 Hz to 100 Hz. Noise associated with the end of blinks and saccades was then removed as follows. For each participant, mean pupil diameter was calculated for every sample during the 6 s time windows following a blink and following a saccade (pupil response). A double gamma impulse response function (IRF) was then fit to the pupil response following a blink. A single gamma IRF was fit to the pupil response following a saccade. The blink and saccade IRFs were convolved with blink and saccade ends to create individually tailored nuisance regressors, and a cleaned data set was then acquired by using the residuals from a linear model describing measured pupil responses as a function of these nuisance regressors. Finally, the previously filtered out slow drift was added back into the data, as this is a meaningful characteristic of pupil size over time.

Tonic pupil size and phasic pupil response were then estimated for each participant and trial of the encoding and detection task. For every trial, tonic pupil size was defined as the mean pupil size in the 500 ms window preceding the trial. In addition, phasic pupil responses were defined as the area under the curve (AUC, using Simpson’s rule; Swallow et al., 2019), where the curve was the double gamma IRF that best fit the pupil time series during the 2 s interval following trial onset. Because AUC is the area between the phasic pupil’s curve following trial onset relative to the pre-trial mean, trials with pupil dilation result in positive AUCs and trials with pupil contractions result in negative AUCs. AUC values were z-scored by subtracting the individual participant’s mean AUC and dividing by the standard deviation to produce the *scaled phasic pupil response*. A linear mixed effects model with random intercepts for participant indicated a significant, negative relationship between the scaled phasic pupil responses and tonic pupil size, β = -.282, 95% CI = [-.320, -.244], *t*(2446) = −14.53*, p* < .001. Residuals from this model were used in subsequent analyses and are referred to as *phasic pupil responses* for simplicity.

The effects of tone type on phasic pupil responses during encoding were evaluated by fitting a linear mixed effects model to the phasic pupil responses (averaged over presentations of an image) with tone type as a fixed effect and random intercepts for image. Random intercepts for participant had near-zero variance and were thus removed from the final model.

#### 2.5.5 Univariate Analysis

Following pre-processing, EPI volumes with motion greater than 0.3 mm were excluded and the data were spatially smoothed using a Gaussian kernel until blur reached a full-width-half-maximum of 5.0 mm (3dBlurtoFWHM). To better estimate activity in the LC, voxels in the neighboring fourth ventricle, labeled with FreeSurfer (recon-all), were excluded from smoothing and subsequent analyses. In addition, masks defining the spatial extent of the brain in the aligned anatomical and EPI data sets, excluding the fourth ventricle, were applied to the EPI data. Data were scaled to a mean of 100 and a range of 0-200 to allow interpretation of beta weights as percent change.

Responses to events of different types were estimated for each voxel using 3dDeconvolve. All models included six motion regressors and 3rd order polynomial drift in baseline as nuisance variables. Regressors of interest were created by convolving a delta function for each event of interest with the two-parameter SPMG2 hemodynamic response function (HRF; Henson et al., 2002). In the univariate encoding and detection task analyses, regressors were included for each combination of tone type and image type, for a total of 18 regressors of interest. When using the SPMG2 HRF, 3dDeconvolve produces two beta estimates for each condition. These were used to estimate the first 5 timepoints (12.5 s) of the hemodynamic response to each of the 18 conditions for subsequent group level analyses.

Univariate analyses of the ROIs were performed by extracting the mean estimated HRF across voxels located within the boundaries of the ROIs for each of the 18 conditions. For each ROI, estimated HRFs were additionally averaged across image type and analyzed in R with a linear mixed effects model that included tone type, time (timepoints 0 s – 12.5 s), hemisphere (left, right), and all interactions as fixed effects and random intercepts for participant and image type. The one exception was the LC ROI, which was collapsed across hemispheres. Models were simplified by excluding the interactions with hemisphere for ROIs that did not show a hemisphere by tone type interaction (all but MC). To characterize the effects of tone type over time in each ROI, general linear tests comparing activity across encoding conditions were then performed for each time point. Although all time points were tested, we focus on time points 2.5 s – 7.5 s in our report. We expected the hemodynamic response to peak within that time frame because the stimuli were brief (cf. Hu et al., 2010) and the ABE, by its nature, should operate quickly.

Whole brain, group-level univariate analyses were performed to characterize the effects of target and distractor tones on activity throughout the brain. Voxels for which there was a significant interaction of tone type and time were identified in an analysis of variance, with tone type, image type, and time (timepoints 0 s – 12.5 s) as factors, using 3dMVM (Chen et al., 2015). To further characterize the interaction of tone type and time, the statistical map for this interaction was thresholded at a False Discovery Rate of *q* < .001 (Genovese et al., 2002) to create a mask of voxels whose hemodynamic response significantly differed across the three tone type conditions. Post hoc paired t-tests (3dttest++) on voxels within the tone type by time interaction mask were performed on timepoints 2.5 s – 7.5 s of the estimated HRF to target vs. distractor tones, target tones vs. no tones, and distractor tones vs. no tones. Statistical and cluster size thresholds were used to correct for multiple comparisons based on simulations that used spatial auto-correlation functions (using AFNI function 3dClustSim; Cox et al., 2017).

#### 2.5.6 Trial-Specific Activity Estimation

Trial-specific activity was estimated by fitting a separate linear model for each trial of the encoding and detection task using the least square-separate approach (Mumford et al., 2012). The deconvolution was performed using AFNI’s 3dDeconvolve with the SPMG2 option, such that each single-trial response was modeled by two regressors (a gamma response function and its time derivative). Similarly, the combined responses on all other trials were modeled by two nuisance regressors. In addition, the design matrix included the same motion and drift nuisance variables used in the univariate model described above. The single-trial gamma function estimates were then saved, resulting in 576 (4 runs x 144 trials) beta maps.

#### 2.5.7 Beta Series Correlation Analysis

The effects of tone type on functional connectivity between the HPC and visual ROIs was estimated using beta series correlations (Rissman et al., 2004; Cisler et al., 2014; Geib et al., 2017) generated from the trial-specific activity estimates. To avoid introducing distortions, we did not subtract the mean pattern from each voxel or scale the data prior to computing these values (Garrido et al., 2013).

First, we concatenated the beta weights of trials sharing the same tone type condition (resulting in three series per voxel with 192 elements each). To obtain ROI-to-ROI functional connectivity estimates, separately for each tone type condition, we generated a mean beta series (obtained by averaging the series across voxels) for each ROI and computed pairwise Fisher-transformed Pearson correlations between those. We fit a linear mixed effects model to the correlation coefficients, with tone type and ROI pair as fixed factors and random intercepts for participant. Holm-corrected general linear tests comparing the different levels of tone type were performed.

ROI-to-voxel beta correlation analyses were then performed for each hippocampal ROI to test the hypothesis that communication should increase between the HPC and visual regions following target tones. Fisher-transformed Pearson correlations between the mean beta series of the seed ROI and each voxel in all visual and hippocampal ROIs (including those in the seed ROI) were then calculated for each tone type and participant. Linear mixed effects models (3dLME) with tone type as a fixed effect and random intercepts for participant were then fit to the ROI-to-voxel correlations. General linear tests contrasted the target with the distractor and no tone conditions at each voxel. Candidate clusters in the resulting maps were identified after accounting for spatial autocorrelation in the data (estimated with 3dFWHMx) and by thresholding based on a minimal cluster size and maximal p-value (voxel edges must touch, *a* = .05, uncorrected *p* = .05; 3dClustSim and 3dClusterize; Cox et al., 2017). This was followed by confirmatory analyses in which each cluster was treated as an ROI. The correlation between the average beta series of the ROI and that of the respective hippocampal seed was computed separately for each participant. A linear mixed effects model, with tone type as a fixed effect and random intercepts for participant, was fit to the correlations. Follow-up general linear tests contrasting the tone type conditions were then performed. This was done to ensure that the clustering procedure did not produce spurious clusters.

#### 2.5.8 Support Vector Classification

On their own, differences in BOLD magnitude do not necessarily indicate changes in the quality or extent of stimulus processing (e.g., Hatfield, McCloskey & Park, 2016; Ward, Chun & Kuhl, 2013). We therefore used linear support vector machine (SVM) classification (Suykens & Vandewalle, 1999; Hsu & Lin, 2002) to probe the effects of target tone detection on image category decoding accuracy in the visual and hippocampal ROIs (V1, V2, FG, PG, aHPC, and pHPC). The algorithm estimates a hyperplane that maximizes the margin between it and samples that belong to different classes (Suykens & Vandewalle, 1999; Hsu & Lin, 2002). It is among the most common approaches to multivoxel pattern analysis (Mahmoudi, Takerkart, Regragui, Boussaoud & Brovelli, 2012; Haxby, Connolly & Guntupalli, 2014; Diedrichsen & Kriegeskorte, 2017). Importantly, it sidesteps many of the interpretability issues inherent to representational similarity analysis (cf. Walther et al., 2016).

Each trial-wise beta map was assigned one of six labels indicating the type of image presented on that trial. Individual beta series maps were standardized across trials such that each voxel had a mean of zero and a standard deviation of one. Classification accuracy was estimated for each tone type and ROI using repeated 4-fold cross validation. Balanced training sets of 144 trial-wise beta maps were randomly drawn 30 times (the remaining 48 trials in each iteration were reserved as a test set); on each iteration a new linear SVM (C=1, one-vs-one multiclass, default implementation in scikit-learn 0.21.2; Pedregosa et al., 2011) was fit to the training set and applied to the test set to obtain a confusion matrix and a classification accuracy estimate for each tone type condition. These were averaged across the 30 iterations to produce one estimate and one confusion matrix for each combination of participant, tone type, and ROI. The effects of tone type on mean classification accuracy were evaluated for each ROI using linear mixed effects models with tone type as a fixed effect and random intercepts for participant.

#### 2.5.9 LC Functional Connectivity

ROIs that exhibited functional connectivity with LC during rest were identified using resting state data reported in Turker et al. (2021). Briefly, intrinsic functional connectivity (iFC) maps were created for each participant using denoised multi-echo data and the participant’s individually defined LC ROI. Data were denoised using ME-ICA (Kundu et al., 2013), bandpass filtered (0.01 < f < 0.1), and were not additionally blurred. A group-level iFC map was created using voxel-wise t-tests (3dttest++) and a one-sided clustering procedure at *p* = .01 and FDR = .018 (3dClusterize; corrected for multiple comparisons using the false discovery rate (FDR = .02; Genovese et al., 2002). Twenty ranked peaks were extracted from the group iFC map (3dmaxima) with a minimal distance of 18 mm (6 voxels) between peaks. Next, 6 mm spherical ROIs were constructed around those peaks and a final set of twenty ROIs was obtained by intersecting the spheres with the group-level iFC map thresholded at *p* < .001, *q* < .004, producing the final LC-iFC ROIs (Table S1 in the *Supplementary Materials*).^1^

### 2.6 Statistical Software

All group-level analyses were performed in R v3.6.1 (R Core Team, 2013) or in AFNI v16.2.07 (using 3dLME or 3dMVM). Unless otherwise noted, linear mixed effects models were fit using lme4 v1.1.21 (Bates et al., 2015). Type III (Satterthwaite’s method) ANOVA tables were obtained using the ‘joint_tests’ function in the package emmeans v1.3.5.1 (Length, 2019). General linear tests were performed and uncorrected confidence intervals were obtained using the emmeans functions ‘contrasts’ and ‘confint.’ In all analyses, Holm-Bonferroni adjusted p-values were computed separately for each set of tone type comparisons. Confidence intervals, where reported, are uncorrected.

## 3. Results

### 3.1 Behavioral Task Performance and Whole Brain Analysis

Participants accurately performed the detection task, pressing the button for M = 97.5% of the targets, SD = .15, 95% CI = [.970, .990], M = 5.1% of the distractors, SD = .22, 95% CI = [.023, .060] and M = 0.2% of the no tone trials, SD = .05, 95% CI = [.001, .003]. Incorrect button presses were more likely to follow a distractor tone than no tone, *t*(20) = −4.16, *p* < .001, *d* = 1.25.

The tone type condition did not significantly influence image recognition accuracy, *F*(2, inf) = 1.08, *p* = .34. However, it did influence the confidence with which images were correctly recognized, *F*(2, 1660.8) = 6.81, *p* = .001. Participants reported higher levels of confidence for correctly recognized images paired with a target than for those paired with a distractor (*M_Diff_* = 5.58), 95% CI = [1.83, 9.32], *t*(1899.08) = 3.57, *p* = .001, *d* = .57, or presented without a tone (*M_Diff_* = 4.10), 95% CI = [0.23, 7.97], *t*(1573.68) = 2.54, *p* = .023, *d* = .44, but confidence did not differ between correctly recognized images presented in the distractor and no tone conditions (*M_Diff_* = 1.48), 95% CI = [-2.38, 5.34], *t*(1578.25) = 0.92, *p* = .359, *d* = .10. Target detection during image encoding thus increased the confidence with which those images were later correctly recognized, and distractor rejection did not significantly interfere with the encoding of a concurrently presented image (Figure 1B).

Whole brain analyses revealed that auditory target detection influenced BOLD activity in regions spanning medial occipital, medial parietal, anterior cingulate, superior temporal, middle frontal, and subcortical areas, including thalamus (time by tone type interaction, *F* > 3.483, *q* < .001; Figure 2A). In many of these regions, activity was initially higher on the target tone trials than in the other two types of trials, though this relationship reversed at subsequent timepoints (Figure 2B, top and middle row). Relative to no tone trials, the response to distractor trials was smaller in magnitude than the response to target trials in most regions (Figure 2B, bottom row).

**Figure 2.**
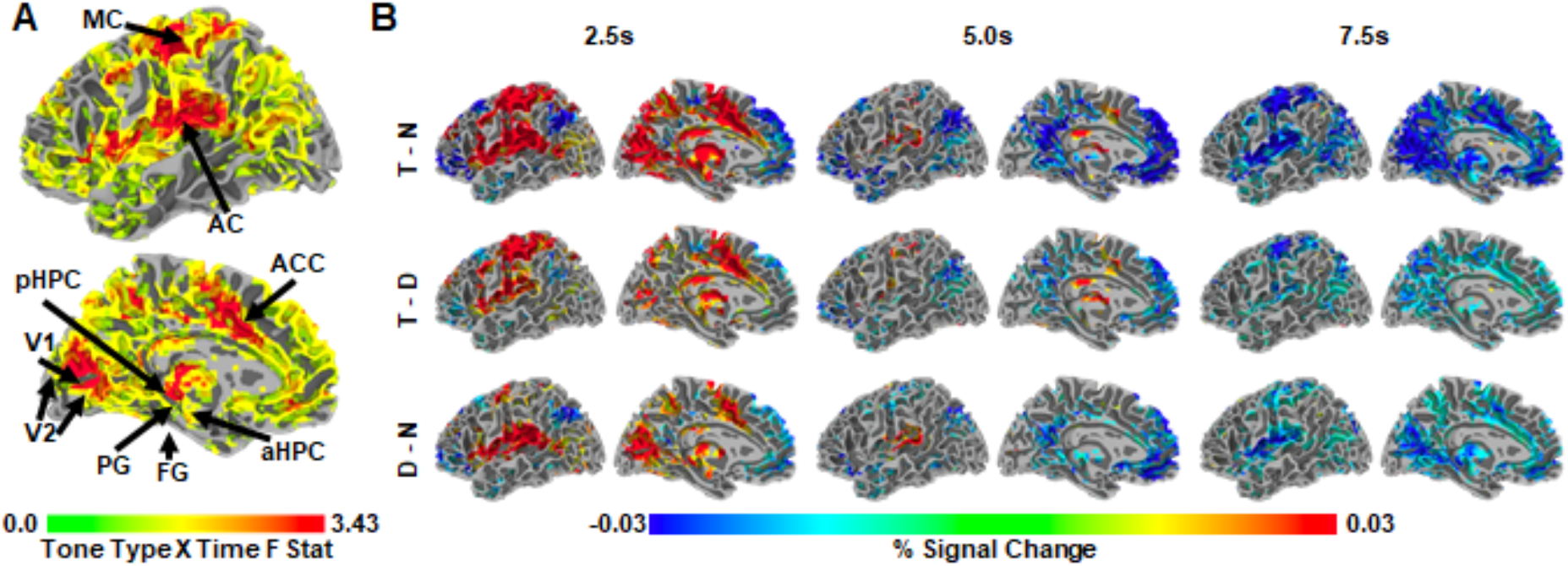
**A**: Whole-brain, group-level F statistic map illustrating the interaction of tone type and time. Only voxels showing a significant tone type by time interaction, F > 3.483, q < .001, in the left hemisphere are shown. Arrows indicate the approximate locations of the ROIs in this study, (see 2.4 in *Materials and Methods;* ROIs were defined for each individual). Note that FG is on the inferior surface and is not visible. Region of interest abbreviations: primary visual cortex (V1), secondary visual cortex (V2), fusiform gyrus (FG), parahippocampal gyrus (PG), posterior hippocampus (pHPC), anterior hippocampus (aHPC), motor cortex (MC), auditory cortex (AC), anterior cingulate cortex (ACC). **B**: Whole-brain, group-level statistical maps illustrating those voxels from (A) that also significantly differed across two tone conditions at 2.5 s, 5.0 s, and 7.5 s after trial onset, z = 1.96, p < .05, range of corresponding q thresholds: .016-.042 for 2.5 s, .049-.111 for 5.0 s, and .02-.06 for 7.5 s. Left to right: time points 2.5 s, 5.0 s, and 7.5 s. Top to bottom: target – no tone baseline (T-N), target – distractor (T-D), distractor – no tone baseline (D-N). All statistical maps are overlaid on the MNI N27 atlas.

### 3.2 Locus Coeruleus and Phasic Pupil Responses

To test our hypothesis that the LC is involved in the ABE, we examined whether target tone trials evoked greater phasic pupil responses and LC signal changes than did distractor tone and no tone trials. Consistent with this possibility, BOLD responses in the hand-traced LC ROIs (Figure 3A) exhibited an interaction of tone type and time, *F*(10, 2225) = 8.28, *p* < .001, reflecting greater increases in activity at 2.5 s on target tone trials than on distractor tone trials, *t*(2225) = 6.51, *p* < .001, *d* = .48, and no tone trials, *t*(2225) = 7.03, *p* < .001, *d* = .58 (Figure 3B). Phasic pupil responses also varied across tone type, *F*(2, 2124.6) = 336.98, *p* < .001: they were more positive on trials that included a target tone than on trials that included a distractor tone (*M_Diff_* = 0.47), 95% CI = [0.37, 0.56], *t*(2110) = 11.45, *p* < .001, *d* = .25, or no tone (*M_Diff_* = 1.05), 95% CI = [0.96, 1.15], *t*(2102) = 25.86, *p* < .001, *d* = .39. They were also greater on distractor tone trials than on no tone trials (*M_Diff_* = 0.59), 95% CI = [0.49, 0.68], *t*(2160) = 14.44, *p* < .001, *d* = .13 (Figure 3C). These data demonstrate increased activity of the LC system on trials that require a response and provide no evidence for inhibitory effects of distractor rejection on this system.

**Figure 3.**
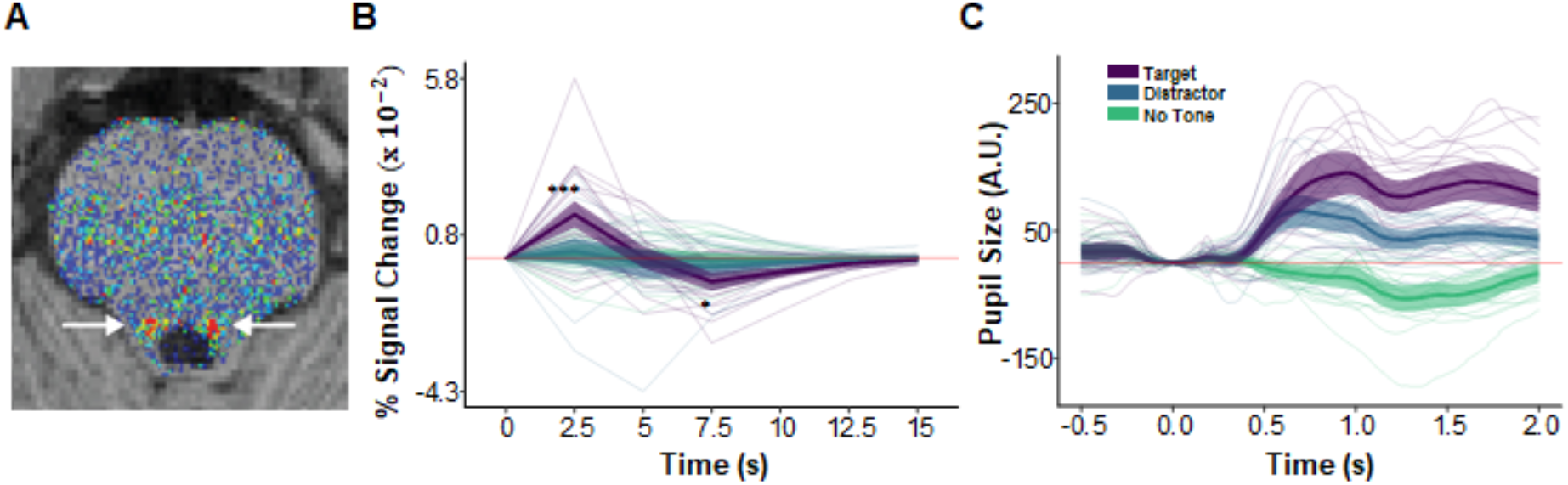
Auditory target detection increased phasic pupil responses and LC activity. **A**: An axial slice of a corrected neuromelanin weighted T1 scan following the procedure in Turker et al. (2021) to individually localize LC (arrows). **B**: BOLD magnitude (% signal change) time series for the LC. Asterisks indicating significant differences between target (purple) and distractor (blue) conditions are shown only for 2.5 s – 7.5 s (*p* < .05*, *p* < .01**, *p* < .001***). **C**: Phasic pupil response magnitude time series during the image encoding and target detection task, as a function of subsequent image recall. In panels B and C faint lines show data for a single participant. Thick lines show the mean across participants and ribbons around the thick lines indicate the 95% confidence intervals around the mean.

### 3.3 BOLD Responses in Perceptual and Motor Regions

To examine the effects of target detection on the processing of episodic information, planned analyses tested whether tone type modulated BOLD magnitude within regions involved in stimulus processing, encoding, and response generation: bilateral MC, V1, V2, FG, PG, aHPC, and pHPC. These analyses evaluate whether previously reported effects of auditory target detection on BOLD responses in visual cortex (Swallow, Makovski & Jiang, 2012) (1) reflect an increase over a neutral baseline condition as well as over distractor conditions, and (2) are present in other regions important for episodic encoding.

The interaction of tone type and time was significant in V1, V2, FG, PG, aHPC, and pHPC, smallest *F*(10, 4492) = 12.48, *p* = .001 for FG. Extending earlier reports (Swallow, Makovski, & Jiang, 2012), V1 showed a larger initial increase in activity on target trials than on both distractor and no tone trials, smallest *z* = 7.81, *p* < .001, *d* = .43. Similar increases were also observed in V2, FG, and PG, smallest *z* = 2.60, *p* = .019, *d* = .15, as well, demonstrating that these effects can also be detected in higher-level visual regions as well as in early visual cortex. Additionally, in all cases BOLD activity showed a steeper drop-off in magnitude on target trials relative to distractor and no tone trials (Figure 4; see also Supplementary Figure S1).

**Figure 4.**
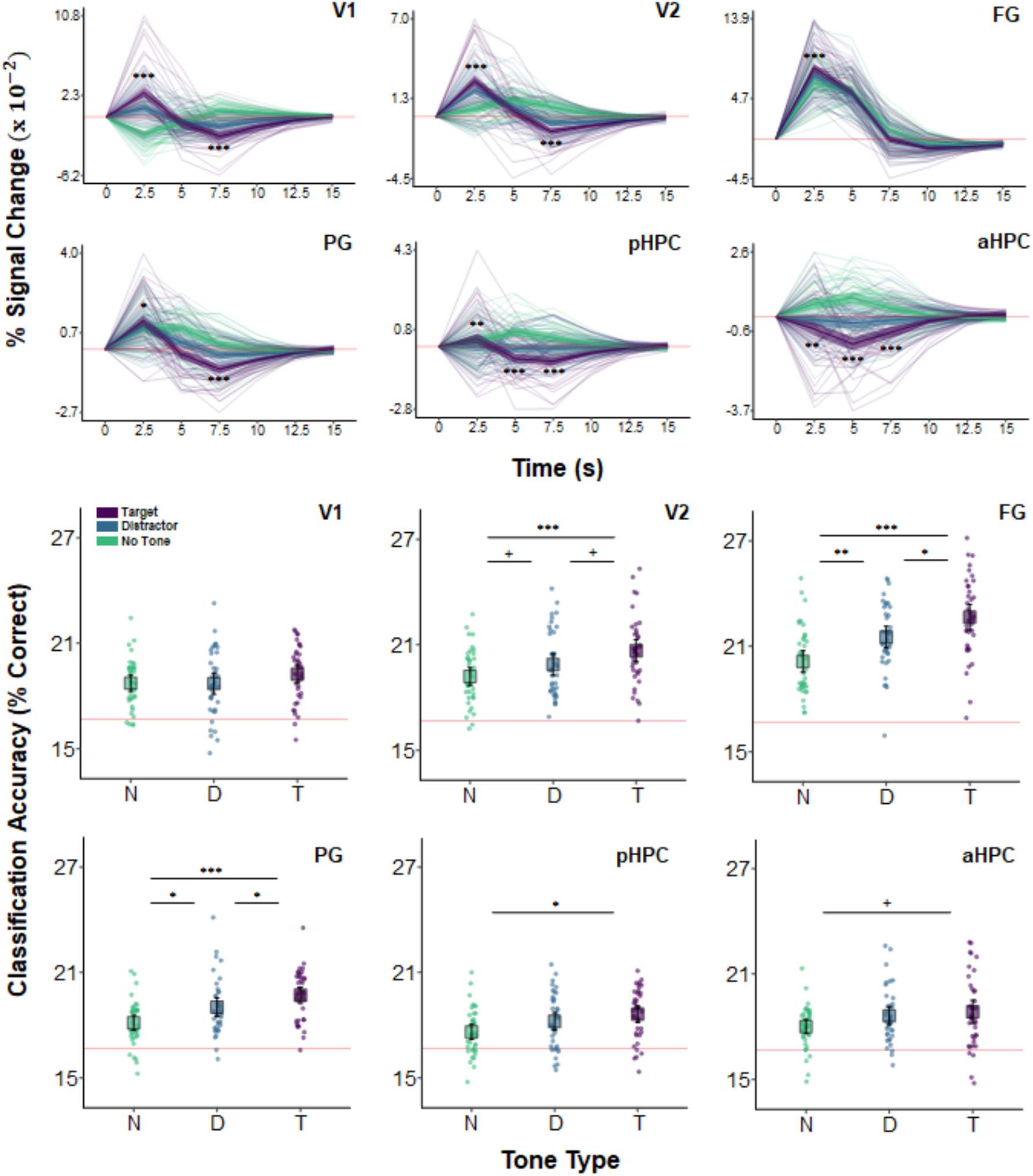
BOLD magnitude and image category classification accuracy as a function of tone type. Top two rows: BOLD magnitude (% signal change) time series for the visual and HPC ROIs in the target (purple), distractor (blue), and no tone (green) conditions. Asterisks indicating significant differences between target and distractor conditions are shown only for 2.5 s – 7.5 s (*p* < .05*, *p* < .01**, *p* < .001***). Faint lines show data for a single participant. Thick lines show the mean across participants. Ribbons around the thick lines indicate the 95% confidence intervals around the mean. Bottom two rows: six-way image category classification accuracy in the same ROIs. Theoretical chance-level performance (16.67%) is marked by a red horizontal line. Each point represents classification accuracy for a single participant. Large squares are centered on the sample mean and error bars indicate 95% confidence intervals around the mean. Asterisks denote a significant difference (*p* < .1+, *p* < .05*, *p* < .01**, *p* < .001***). Region of interest abbreviations: primary visual cortex (V1), secondary visual cortex (V2), fusiform gyrus (FG), parahippocampal gyrus (PG), posterior hippocampus (pHPC), anterior hippocampus (aHPC), motor cortex (MC), auditory cortex (AC). See 2.4 in *Materials and Methods* for ROI definitions.

The HPC generally showed larger decreases in activity on target trials than on distractor and no tone trials. This was true of both the aHPC and pHPC, which at 5 s were more strongly deactivated on target trials than on distractor trials, smallest *z* = 4.50, *p* = .001, *d* = .28, and no tone trials, smallest *z* = 11.81, *p* < .001, *d* = .751. However, activity in the aHPC decreased more rapidly than it did in the pHPC: at 2.5 s, activity in the aHPC was lower on target than on distractor, *z* = 3.12, *p* = .002, *d* = .18, and no tone, *z* = 8.54, *p* < .001, *d* = .52, trials, whereas activity in pHPC was higher on target trials than on distractor trials, *z* = 3.49, *p* = .001, *d* = .20 (Figure 4).

Except for in the MC, there were no interactions between hemisphere and tone type, largest *F*(2, 4475) = 1.80, *p* = .165, or between tone type, time, and hemisphere, largest *F*(10, 4475) = 1.77, *p* = .061. In MC, at 2.5 s, BOLD activity was greater on target trials than on distractor and no tone trials in the left hemisphere, smallest *t*(2225) = 13.55, *p* < .001, *d* = .98.

These results demonstrate that target detection modulated the magnitude of activity in regions involved in representing visual and episodic information.

### 3.4 Decodable Stimulus Information in the Visual Cortex and Hippocampus

Increased BOLD activity on target trials does not necessarily entail that processing in these regions is enhanced. To test our hypothesis that target detection facilitates the processing of concurrently presented images, we conducted analyses of image category classification accuracy using voxel-wise patterns of activity in the visual and hippocampal ROIs. These revealed a main effect of tone type in V2, FG, PG, and pHPC, smallest *F*(2, 92) = 4.59, *p* = .013 for pHPC, but not in V1 or aHPC, largest *F*(2, 92) = 1.36, *p* = .261 for V1. Follow up analyses indicated that classification accuracy was higher on target trials than on no tone trials in V2, FG, PG, and pHPC, smallest *t*(92) = 3.01, *p* = .01, *d* = .96 for pHPC. Classification accuracy was also higher on target compared to distractor tone trials in FG *t*(92) = 2.60, *p* = .011, *d* = .62, PG, *t*(92) = 2.09, *p* = .039, *d* = .52, and marginally in V2, *t*(92) = 2.17, *p* = .065, *d* = .44. Similarly, relative to no tone trials, distractor trials significantly enhanced classification accuracy in FG and PG, smallest *t*(92) = 2.764, *p* = .014, *d* = .87 for the latter, and showed a marginal effect in V2, *t*(92) = 1.88, *p* = .065, *d* = .43 (Figure 4). Thus, regions whose activity was enhanced by target detection also exhibited within-ROI patterns of activity that better correlated with image category on these trials than on distractor or no tone trials. This was particularly true in higher-level visual areas, FG and PG, which should be more tuned to image categories than V1 and V2.

### 3.5 Visuo-Hippocampal Functional Connectivity

To test our hypothesis that target detection enhances communication between regions involved in episodic processing, functional connectivity between all visual and hippocampal ROIs was quantified with beta series correlations (Figure 5). Our analysis examined the effect of tone type across each pairing of the following ROIs: l/r-V1, l/r-V2, l/r-FG, l/r-PG, l/r-aHPC, and l/r-pHPC. Tone type and ROI pair influenced functional connectivity, *F*(2, inf) = 46.65, *p* < .001 and *F*(77, inf) = 122.24, *p* < .001, respectively, but did not interact, *F*(154, inf) = 0.33, *p* > .999. Holm-corrected general linear tests indicated that functional connectivity was enhanced on target trials relative to both distractor trials, *z* = 9.48, *p* < .001, *d* = .50, and no tone trials, *z* = 6.36, *p* < .001, *d* = .32. Functional connectivity was also higher on no tone trials than on distractor tone trials, *z* = 3.12, *p* = .002, *d* = .24. Thus, relative to the no tone trials, target tones increased correlations between ROIs while distractor tones decreased them.

**Figure 5.**
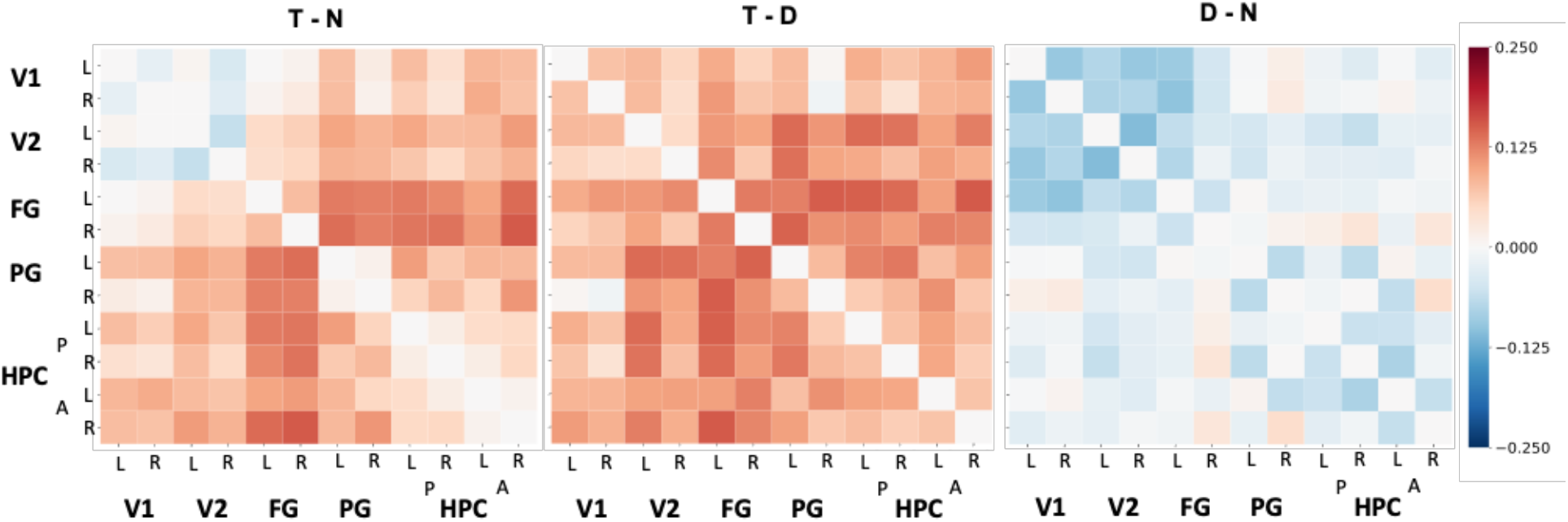
Differences in functional connectivity throughout the visual cortex and HPC by tone type. Group-level, pairwise functional connectivity differences across tone type conditions (left to right: target – no tone baseline, target – distractor, distractor – no tone). Each cell corresponds to the raw difference between the Fisher-transformed beta series correlations of the anatomical ROIs specified in the axis labels (L = left, R = right, A = anterior, P = posterior) in the two conditions compared in its respective matrix. A linear mixed effects model and general linear tests indicated that beta series correlations were significantly greater for target tones than distractor tones, but that this effect did not interact with ROI pair.

To more precisely identify the regions whose functional connectivity with the HPC changed with tone type, we calculated ROI-to-voxel functional connectivity maps and extracted candidate clusters by contrasting the tone type conditions. We refer to these clusters by the ROI seed that generated them and the anatomical area that they overlapped with most (e.g., l-pHPC<->r-V2 refers to a cluster largely overlapping with r-V2 that was correlated with l-pHPC). No clusters were found when the distractor condition was contrasted with the no tone baseline, indicating that the distractor tones did not reliably alter functional connectivity between the HPC and visual cortex. However, three clusters (l-pHPC<->l-FG, l-pHPC<->r-FG, and r-aHPC<->l-FG) were identified when target trials were contrasted with no tone trials and seven clusters were identified when target trials were contrasted with distractor trials (Table 1 and Figure 6). Both sets of clusters spanned voxels throughout visual cortex.

**Figure 6.**
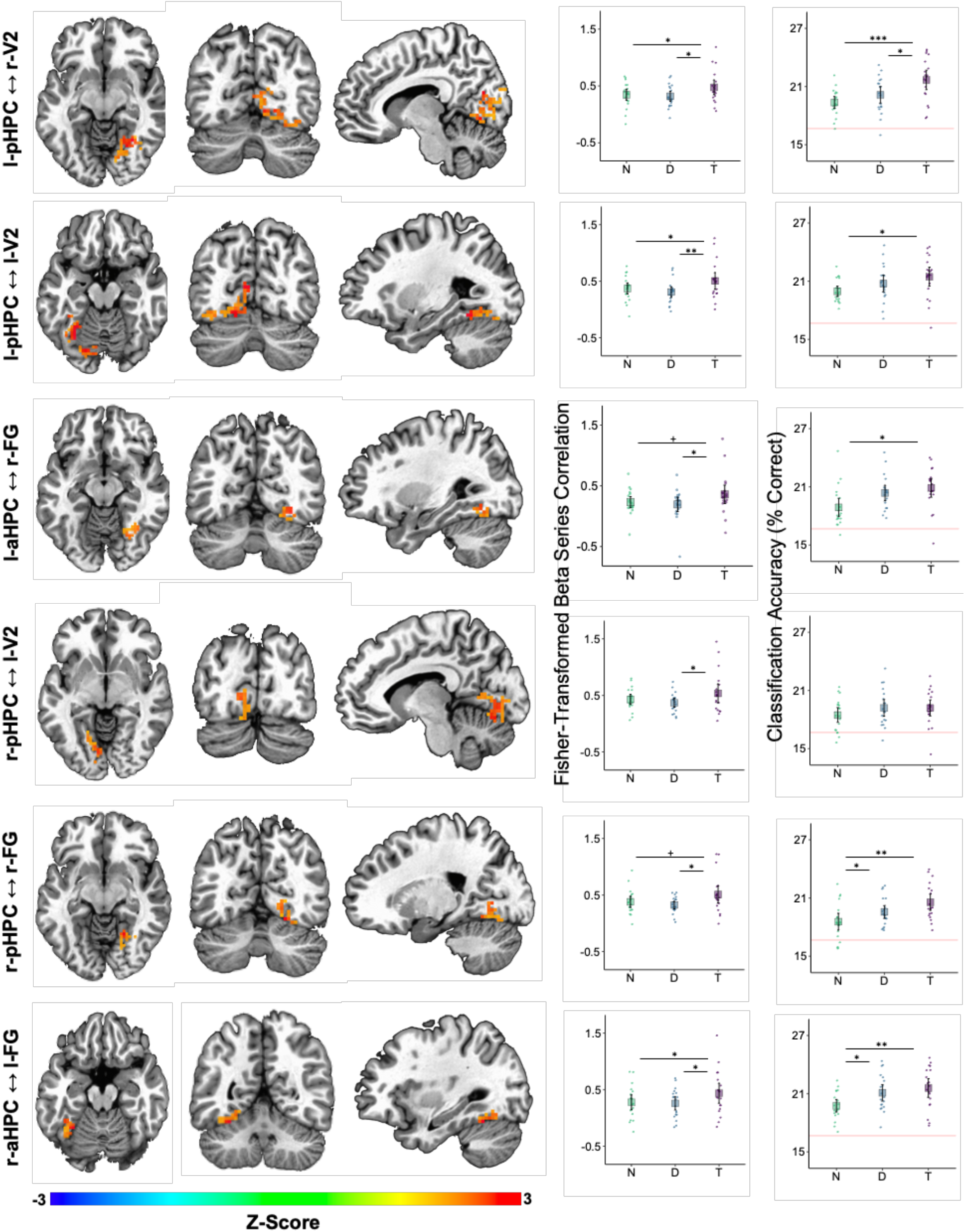
Auditory target detection improved image classification accuracy in visual regions showing increased functional connectivity with the HPC. Each row corresponds to one cluster detected as being more strongly correlated with the HPC on target tone trials than on distractor tone trials. The seed-cluster pair label is denoted on the left. Left three columns: Voxel colors correspond to z-scores obtained by contrasting the Fisher-transformed correlations between the mean beta series for the hippocampal seed region and that of each voxel in V1, V2, FG, PG, pHPC, and aHPC. Middle column: mean Fisher-transformed beta series correlations between each cluster and its respective hippocampal seed, obtained separately for each tone type condition—no tone (N), distractor tone (D), and target tone (T). Right column: six-way image category classification accuracy for each cluster. Theoretical chance-level performance (16.67%) is marked by a red horizontal line. Middle and Right columns: large squares are centered on the sample mean and error bars indicate 95% confidence intervals around the mean. Each point represents an observation from a single participant. Asterisks denote a significant difference (*p* < .05*, *p* < .01**, *p* < .001***).

**Table 1.** Clusters showing higher functional connectivity with the HPC on target tone than on distractor tone trials in the seed-to-voxel beta series correlation analysis.

| Seed | Cluster | Center of Mass | Size | % Overlap with Anatomical ROIs |  |  |  |  |  |  |  |
| --- | --- | --- | --- | --- | --- | --- | --- | --- | --- | --- | --- |
|  |  |  |  | l-V1 | r-V1 | l-V2 | r-V2 | l-FG | r-FG | l-PG | r-PG |
| l-pHPC | r-V2 | -16.0, 72.5, -1.1 | 264 | 0.8% | 23.6% | 1% | 50.2% | - | 24.4% | - | - |
| l-pHPC | l-V2 | 20.3, 69.0, -10.8 | 253 | 8.3% | - | 48.8% | - | 42.9% | - | - | - |
| l-aHPC | l-FG | 32.4, 52.6, -17.9 | 60 | - | - | - | - | 100% | - | - | - |
| l-aHPC | r-FG | -30.5, 61.9, -13.9 | 43 | - | - | - | 10.5% | - | 89.5% | - | - |
| r-pHPC | l-V2 | 10.8, 74.1, -5.4 | 102 | 17.5% | - | 79% | - | 3.5% | - | - | - |
| r-pHPC | r-FG | -22.9, 64.3, -10.7 | 56 | - | 1.2% | - | 48.9% | - | 49.9% | - | - |
| r-aHPC | l-FG | 25.1, 66.4, -14.5 | 148 | 1% | - | 37.5% | - | 61.5% | - | - | - |

**Note:** Center of mass is reported in Right Anterior Inferior coordinates (RAI, the AFNI default), volumes are in mm<sup>3</sup>, and the percentages of voxels in a cluster that overlapped with each anatomical ROI are the mean across subjects.

Confirmatory analyses on clusters identified in the target versus distractor contrast were performed by averaging functional connectivity across all voxels within a cluster and then testing the effect of tone type in a linear mixed effects model. This analysis indicated that all pairs showed an effect of tone type on functional connectivity, smallest *F*(2, 36) = 3.77, *p* = .033 for l-aHPC<->r-FG, except l-aHPC<->l-FG, *F*(2, 36) = 3.02, *p* = .061. In those pairs showing an effect of tone type, functional connectivity was higher on target trials than on distractor trials, smallest *t*(36) = 2.61, *p* = .039, *d* = .54 for l-aHPC<->r-FG. Functional connectivity was also higher on target trials than on no tone trials in the pairs l-pHPC<->r-V2, l-pHPC<->l-V2, and r-aHPC<->l-FG, smallest *t*(36) = 2.39, *p* = .045, *d* = .46 for l-pHPC<->l-V2. No differences in functional connectivity were found between the distractor and no tone trials in any of the clusters, largest *t*(36) = 1.12, *p* = .273, *d* = .29 (Figure 6).

To test the hypothesis that increased visuo-hippocampal coordination during target tone trials is associated with better visual processing, we treated the ROI-to-voxel functional connectivity clusters as ROIs in an image category classification analysis. Tone type affected classification accuracy in all clusters except r-pHPC<->l-V2, smallest *F*(2, 36) = 3.53, *p* = .04. This effect reflected greater accuracy on target trials than on no tone trials, smallest *t*(36) = 2.65, *p* = .035, *d* = .86. Only the l-pHPC<->r-V2 cluster showed higher classification accuracy on target trials than on distractor trials, *t*(36) = 2.54, *p* = .031, *d* = .74. Accuracy was higher on distractor trials than on no tone trials in r-pHPC<->r-FG and r-aHPC<->l-FG, smallest *t*(36) = 2.48, *p* = .036, *d* = .77 for the former (Figure 6).

### 3.6 Functional Connectivity of Regions Associated with the LC

A functional connectivity analysis with the LC as a seed revealed small regions within l-FG, r-FG, and r-HPC that showed higher functional connectivity with the LC in the target tone condition than in the distractor tone condition (respectively, *p*s > .007, *p*s > .012, *p*s > .045; illustrated in Supplementary Figure S4). However, these regions were not large enough to survive corrections for multiple comparison using cluster size 32 (*a* < .05). Voxels in l-MC also showed no evidence of differential connectivity with the LC on target relative to distractor trials, suggesting that this analysis may not have been powerful enough to detect differences in LC connectivity across conditions.

However, if the LC influences activity in regions involved in episodic encoding, then functional connectivity between HPC and regions whose activity is modulated by the LC during rest should be greater on target trials (when LC activity is strongest) than on distractor and no tone trials. To test this hypothesis, a set of 20 regions whose activity was associated with LC activity during a separate resting state scan—referred to as LC-iFC ROIs—were identified (see 2.4 in *Materials and Methods* and Turker et al., 2021; Supplementary Table S1) and their functional connectivity to the four hippocampal seeds was examined. A separate model was fit for each of l-pHPC, r-pHPC, l-aHPC, and r-aHPC.

Main effects were found for tone type, smallest *F*(2, 1062) = 10.11, *p* < .001 for l-aHPC, and region, smallest *F*(19, 1062) = 13.05, *p* < .001 for r-pHPC, but the two did not interact, largest *F*(38, 1062) = 0.49, *p* = .996 for l-pHPC. General linear tests that collapsed across LC-iFC ROIs showed higher functional connectivity on target trials than on distractor trials, smallest *t*(1062) = 4.45, *p* < .001, *d* = .41 for l-aHPC, and no tone trials, smallest *t*(1062) = 2.45, *p* = .029, *d* = .24 for l-pHPC. Functional connectivity to l-pHPC was lower on distractor trials than on no tone trials, *t*(1062) = 2.06, *p* = .039, *d* = .32. Tone type did not significantly influence functional connectivity between the LC and the LC-iFC ROIs, *F*(2, 1062) = .093, *p* = .911, consistent with the possibility that functional connectivity analyses of LC during the encoding task were not sufficiently powerful.

## 4. Discussion

In this fMRI study we investigated the effects of target detection on how visual stimuli are processed and encoded by the brain. We examined how visual regions, the HPC, and the LC respond to images presented concurrently with an auditory target tone (requiring a motor response), a distractor tone (requiring no response), or no tone. The inclusion of a no tone baseline condition, which was absent in previous fMRI studies of similar effects, allowed us to test for both target-induced enhancement and distractor-induced disruption of encoding, perceptual processing, and functional connectivity. We found that, relative to both other conditions, target tones enhanced image recognition confidence (the ABE), the magnitude of phasic pupil responses, activity in LC and in visual regions, visuo-hippocampal functional connectivity, and image category classification accuracy from multivoxel patterns in FG and PG. Combined, these results suggest that auditory target detection enhances the processing of visual information by increasing inter-areal communication and enhancing the specificity of visual representations.

Consistent with existing evidence suggesting that target detection enhances episodic encoding in the attentional boost effect (Swallow & Jiang, 2013), recognition confidence ratings for correctly remembered images were greater when those were paired with a target tone as opposed to a distractor tone. Indeed, previous work has shown that target detection can enhance both recollection and familiarity of images presented during the encoding task even when they are presented one time (Broitman & Swallow, 2020). The absence of significant differences in recognition accuracy in the full sample is consistent with an earlier MRI study (Swallow et al., 2012) and may be a consequence of the relatively long inter-trial intervals that had to be used in this study (cf. Mulligan & Spataro, 2015).

Throughout the visual areas examined, activity in the target tone condition was higher relative to the distractor and no-tone baseline conditions. This pattern was also present in other regions involved in orienting to relevant stimuli. Those included the thalamus, whose higher-order nuclei play a critical role in attention and in the maintenance of sensory representations in awareness and working memory (Saalmann & Kastner, 2011), and the LC, whose noradrenergic projections regulate gain throughout the cortex (Aston-Jones & Cohen, 2005; Servan-Schreiber et al., 1990) and which tends to exhibit larger responses following salient events (Bouret & Sara, 2005). Interestingly, the opposite pattern was observed in aHPC and pHPC: activity decreased on target tone trials relative to the two other conditions. Because the images were presented four times, these decreases may reflect greater repetition suppression of hippocampal activity for target-paired images than for other images (Henson & Rugg, 2003; Summerfield et al., 2008; Larsson & Smith, 2011; Kim et al., 2020). Measures of repetition suppression tend to positively correlate with subsequent recognition memory (e.g., Pihlajamaki et al., 2010) and with functional connectivity between the HPC, visual cortex, and prefrontal cortex (Zweynert et al., 2011). Though our results, as a whole, hint at a relationship between hippocampal repetition suppression and the effects of target detection on recollection, additional work is needed to confirm this possibility.

Changes in inter-areal coordination are thought to play a role in the maintenance and encoding of neural representations (Singer, 2013; Fries, 2005, 2015; Bonnefond et al., 2017; Moyal & Edelman, 2019; Moyal et al., 2020). Enhanced cortico-hippocampal functional connectivity, in particular, has been associated with working memory maintenance and long-term memory encoding (e.g., Gazzaley et al., 2004; for a review, see Poch & Campo, 2012). Target detection could influence perception and memory in a similar fashion, by enhancing communication between perceptual and medial temporal regions in critical moments (e.g., when responding to targets). Our findings are compatible with this idea: functional connectivity between aHPC and pHPC and the visual cortex was higher on target tone trials relative to both distractor and no tone trials. The effect of target tones on HPC to visual cortex functional connectivity was most pronounced for the left pHPC, which showed widespread, bilateral increases in functional connectivity with clusters in the ventral visual cortex—V1, V2, and FG. The aHPC exhibited a similar correlation with FG, but less so with V1 and V2. Differences in the effects of target detection on aHPC and pHPC connectivity are consistent with the differential functional connectivity of the anterior and posterior HPC previously reported in humans (Poppenk et al., 2013; Frank et al., 2019). They also suggest that future research should examine whether target detection has larger effects on the types of information processed by pHPC (perceptual features of stimuli) than on the types of information supported by aHPC (categories of stimuli).

The target-related enhancement of visuo-hippocampal connectivity was accompanied by a boost in image category classification accuracy throughout the ventral visual stream and in pHPC relative to the baseline condition. A similar enhancement was found relative to the distractor tone condition in FG and PG. This suggests that the effect of target detection on subsequent recognition confidence could reflect improved perceptual encoding in higher level visual areas. Surprisingly, a smaller increase in classification accuracy was also observed on distractor tone trials relative to the no-tone baseline in FG and PG. Thus, although exposure to any auditory tone in this task may enhance the quality of visual information processing, target detection provides an additional boost. A similar pattern was also found in smaller ventral visual clusters that exhibited greater functional connectivity with HPC on target compared to distractor tone trials. This suggests a possible relationship between these two effects. Future studies may address the question of whether shifts in long-range coordination are directly related to the quality and extent of perceptual processing and memory encoding (Moyal & Edelman, 2019).

Phasic LC responses have been hypothesized to facilitate the updating of representations and contribute to the ABE (Swallow & Jiang, 2010; Swallow et al., 2012) by enhancing perceptual processing following target detection independent of modality or spatial location (Swallow & Jiang, 2013; Swallow & Jiang, 2014; see also Nieuwenhuis et al., 2005; Bouret & Sara, 2005). Our results are consistent with this view. They suggest a role for the LC in mediating the effects of target detection on perception and memory via the strengthening or reorganization of functional networks. In this study, both LC activity and phasic pupil responses (which may correlate; Joshi et al., 2016) increased on target tone trials relative to both distractor and no tone trials. These differences were observed despite the fact that each condition was equally likely. However, whereas LC activity did not increase on distractor trials relative to no tone trials, pupil diameter did. Indeed, other cognitive and neural factors contribute to pupil size in addition to LC activity (e.g., cognitive effort; Van der Wel & Van Steenbergen, 2018). Additionally, regions that were highly correlated with the LC during rest were also more strongly correlated with aHPC and pHPC on target trials than on distractor trials during the encoding task. This hints at a possible link between target-related LC responses and enhanced hippocampal functional connectivity, which can be addressed directly in future studies.

In the encoding and detection task, target detection differs from distractor rejection in both cognitive and motor demands. However, the effects of target detection on activity in visual cortex, pupil responses, and memory can occur in the absence of an overt motor response and are absent when motor responses are self-generated (Jack et al., 2006; Mulligan, et al., 2016; Swallow & Jiang, 2012; Swallow et al., 2012; Makovski et al., 2013; Swallow et al., 2019; Toh & Lee, 2022). In this study behavioral inhibition in the distractor tone condition also did not lower BOLD magnitude or classification accuracy relative to the baseline, arguing against the possibility of a disruptive effect of response inhibition on processing (as in inhibition-induced forgetting, which has been tied to fluctuations in ventrolateral PFC activity, which were not found here; Chiu & Egner, 2015). While this confirms that our findings reflect a target-induced boost (Swallow & Jiang, 2014), target detection was confounded with motor responses in this study. Therefore, additional research is necessary to identify and confirm the source of the physiological effects we report here, whether it is from the motor system or otherwise (see *Supplementary Materials* for additional analyses).

Though the peak BOLD responses we observed were often earlier than is typical in human fMRI, they are consistent with expected BOLD response latencies (Miezin et al., 2000) and fall within the range of peak latencies reported in investigations of hemodynamic response variability (Handwerker et al., 2004) and BOLD responses to brief (< 1 s) auditory tones (Hu et al., 2010). Rapid BOLD dynamics in our study may also reflect the relatively coarse temporal resolution of the EPI sequence (TR = 2.5 s), as well as the duration and periodicity of stimulus presentation in our task: BOLD responses are sharper and faster when stimuli are brief (Huettel et al., 2004; Hirano et al., 2011; Tian et al., 2010) and both effects may be exaggerated when brief stimuli are presented over a background of periodic stimulation (Lewis et al., 2016). Animal neuroimaging and physiological modeling further suggest that rapid increases and decreases in BOLD signal reflect changes in blood flow and volume within microvasculature supporting temporally and spatially localized neuronal activity (Hirano et al., 2011; Polimeni & Lewis, 2021; Tian et al., 2010). The rapid BOLD dynamics we observed therefore may reflect temporally and spatially precise neural responses to brief stimuli presented during periodic visual change.

Taken together, our data provide novel evidence that target detection facilitates the processing and encoding of information presented at the same time as stimuli that require a response. These events enhance perceptual processing and functional connectivity between the HPC and the ventral visual cortex. Both effects could be related to the stronger LC responses observed following auditory target detection. Our results and interpretation are compatible with the emerging view that, by increasing gain throughout the thalamocortical network at opportune moments, the phasic release of NE from the LC may facilitate functional network reorganization and promote more integrated, information-rich dynamics. Theoretical models and empirical findings have linked higher gain to increases in the topological complexity and variability of population activity (Shine et al., 2018; Moyal & Edelman, 2019) as well as to enhanced inter-regional information transfer (Li et al., 2019). Though LC-mediated changes in gain may be sufficient for mediating the facilitatory effects of target detection on memory, prefrontal influences are also likely to contribute—either by modulating LC output (Jodoj et al., 1998) or by directly regulating hippocampal activity and functional connectivity to support memory encoding (Ranganath et al., 2005; Schott et al., 2011). Future work may combine functional neuroimaging, electrophysiology, and computer simulations to explore these possibilities and provide a precise account of the mechanisms underlying the ABE. This work can further clarify the effects of attending to behaviorally relevant moments on neural dynamics, information representation, and incidental encoding.

## Abbreviations

AC: auditory cortex
ACC: anterior cingulate cortex
AUC: area under the curve
FG: fusiform gyrus
aHPC: anterior hippocampus
pHPC: posterior hippocampus
iFC: intrinsic functional connectivity
IRF: impulse response function;
LC: locus coeruleus
MC: motor cortex
ME: multi-echo
ME-ICA: multi-echo independent component analysis
NE: norepinephrine
nmT1: neuromelanin-weighted T1
PG: parahippocampal gyrus
SVM: support vector machine
V1: primary visual cortex
V2: secondary visual cortex.

## 5. Acknowledgements

The authors would like to thank Suyash Bhogawar, Bohan Li, Alyssa Phelps, Roy Proper, and Emily Qualls for their assistance with data collection. We also thank Elizabeth Riley for her help with identification of the locus coeruleus, and Adam Anderson, Eve DeRosa, and Nathan Spreng for discussion of this work.

## 8. Data Availability Statement

The software used to analyze the data is publicly available and references are provided in the methods. Additional materials are available from the corresponding author upon reasonable request and with proper approval from relevant research and ethics entities. Participants did not provide consent to make these data publicly available.

## Footnotes

1 Though the regions are similar, the exact coordinates and rank order of ROIs in Table S1 differ from those reported in Turker et al. (2021) because of a difference in coordinate systems and a change in how data were compressed during preprocessing.

