## Supplementary Material for "Auditory Target Detection Enhances Visual Processing and Hippocampal Functional Connectivity"

Roy Moyal[1], Hamid B. Turker[2], Wen-Ming Luh[3], & Khena M. Swallow[4] *

1. Department of Psychology & Cognitive Science Program, Cornell University, Ithaca, NY
Postal Address: 211 Uris Hall, Ithaca, NY, 14853, USA


2. Department of Psychology & Cognitive Science Program, Cornell University, Ithaca, New York
Postal Address: 211 Uris Hall, Ithaca, NY, 14853, USA


3. National Institute on Aging, National Institutes of Health, Baltimore, Maryland
Postal Address: 3001 S Hanover St, Baltimore, MD, 21225, USA


4. Department of Psychology & Cognitive Science Program, Cornell University, Ithaca, New York
Postal Address: 211 Uris Hall, Ithaca, NY, 14853, USA


### Activation in Auditory and Motor Cortex

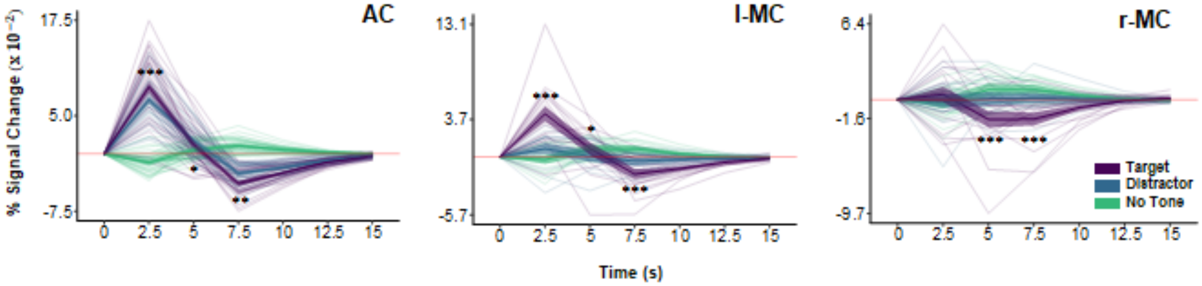


**Figure S1**. BOLD magnitude (% signal change) time series for auditory cortex (AC) and left and right motor cortex (l/r-MC) in the target (purple), distractor (blue), and no tone (green) conditions. Asterisks indicating significant differences between target and distractor conditions are shown only for 2.5 s – 7.5 s (*p* < .05*, *p* < .01**, *p* < .001***). Faint lines show data for a single participant. Thick lines show the mean across participants. Ribbons around the thick lines indicate the 95% confidence intervals around the mean.

### Potential Sources of Modulation

*Anterior Cingulate Cortex* (ACC). To examine potential prefrontal contributions to the behavioral and physiological effects we observed, we performed ROI-to-voxel and ROI-to-ROI beta series connectivity analyses with ACC as a seed and aHPC, pHPC, AC, and l-MC as target ROIs (the procedure was otherwise identical to that used to generate the results described in section 3.5, *Visuo-Hippocampal Functional Connectivity*).

Tone type affected functional connectivity between the ACC and pHPC, $F(2, 88) = 3.90$, $p = .024$, with a target tone advantage over distractor tones, $t(88) = 2.71$, $p = .024$, $d = .53$.

The ROI-to-voxel analysis yielded two significant clusters (presented in Figure S2): ACC<->r-pHPC, center of mass (-25.2, 32.2, -6.9), and ACC<->l-MC, center of mass (40.3, 9.5, 57.8). Both clusters survived a confirmatory functional connectivity analysis, with a main effect of tone type, $F(2, 36) > 3.83$, $p < .031$, and higher correlations in the target condition than in the distractor condition, $t(36) > 2.71$, $p < .031$. Several other clusters, including l-HPC, were too small to survive the corrected significance threshold of 17 voxels ($p < .05$, $a < .05$).

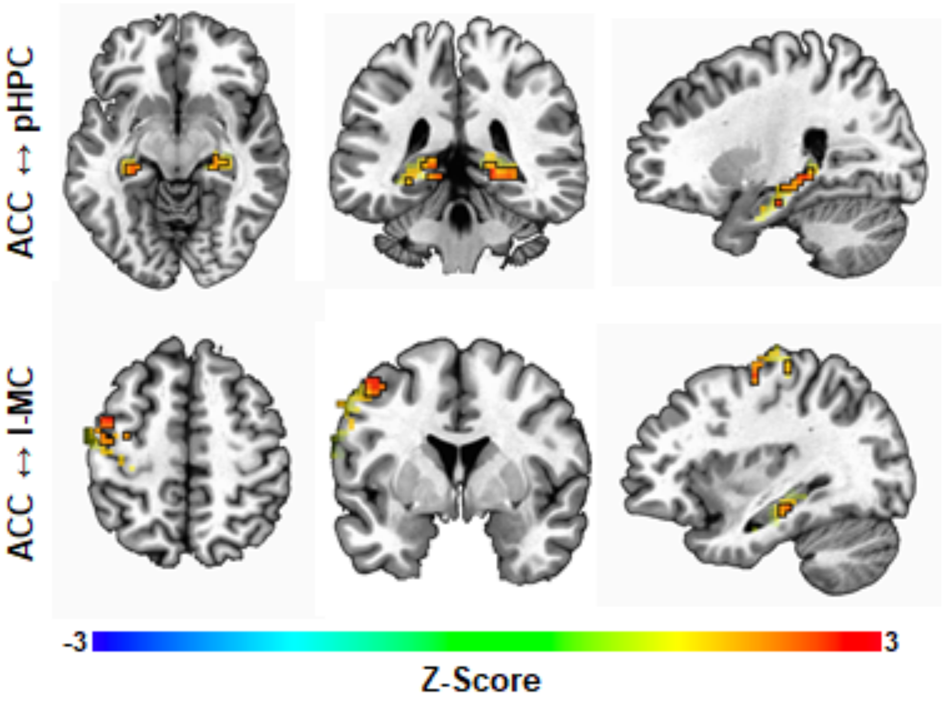


**Figure S2. Anterior Cingulate Cortex Functional Connectivity.** Voxel colors correspond to z-scores obtained by contrasting the Fisher-transformed correlations between the mean beta series for the ACC seed region and that of each voxel in pHPC, aHPC, l-MC, and AC. Voxels that had higher beta series correlations with the ACC on target than on distractor tone trials (uncorrected $p < .05$) are surrounded by a black frame, and other voxels are faded. Each row focuses on one cluster, denoted by the label on the left.

*Locus Coeruleus* (LC). We also performed a beta series connectivity analysis with LC as a seed (whose position and localization procedure are illustrated in Figure S3) and V1, V2, FG, PG, pHPC, and aHPC as target ROIs. Though we found no significant clusters, this ROI-to-voxel analysis yielded several regions within the visual cortex and hippocampus that were more correlated with LC after target tones relative to distractor tones at a threshold of $p < .05$ (Figure S4).

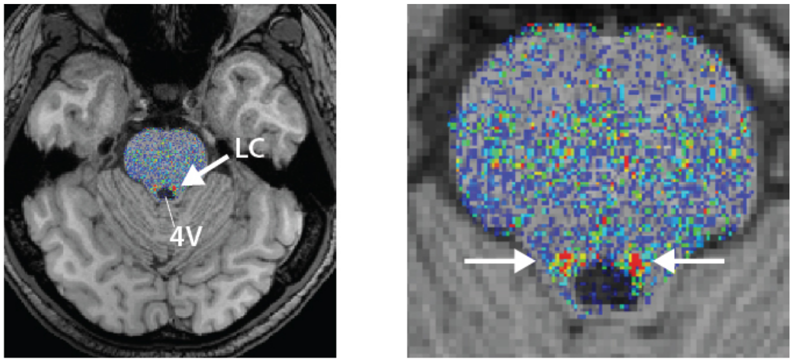


**Figure S3. A visualization of the LC localization process**. Individual MPRAGE scans (including skull) were aligned to the normalized T1-weighted neuromelanin scan to preserve the higher in-plane spatial resolution of the nmT1 images. After extracting the brainstem from the nmT1, correcting image intensity, and setting the false color palette to the predefined range (low values in blue, high values in red, range = 10–80), potential LC voxels can now be visually distinguished from nearby regions (left panel). Two researchers independently traced the LC according to the protocol described in the methods, by selecting voxels that appeared yellow or redder (corresponding roughly to intensity values ≥ 40; see Turker et al., 2021 for more information).

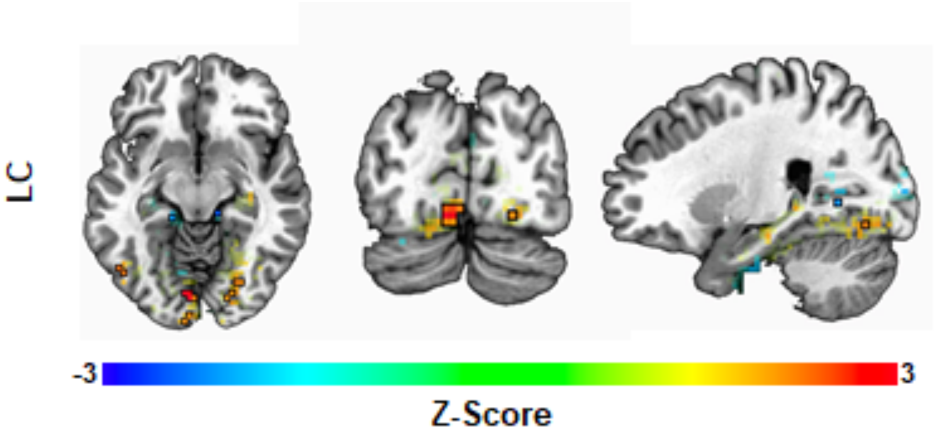


**Figure S4. Locus Coeruleus Functional Connectivity.** Voxel colors correspond to z-scores obtained by contrasting the Fisher-transformed correlations between the mean beta series for the LC seed region and that of each voxel in V1, V2, FG, PG, pHPC, aHPC, l-MC, and AC. Voxels detected as being more strongly correlated with the LC on target tone trials than on distractor tone trials (uncorrected $p < .05$) are surrounded by a black frame, and other voxels are faded.

**Locus coeruleus intrinsic functional connectivity clusters**

**Table S1.** RAI coordinates and sizes ($mm^3$) of the clusters obtained by intersecting the thresholded resting state LC iFC map, $p < .001$, with non-overlapping spherical ROIs centered on the 20 highest z-scores.

| ID | Center of Mass (RAI) | Size ($mm^3$) | Area |
|---|---|---|---|
| 1 | 1.3, 66.7, 12.0 | 837 | L. lingual gyrus / parieto-occip. fissure |
| 2 | -1.8, 27.9, -26.8 | 783 | Pons |
| 3 | -28, -53.1, 17.7 | 729 | R. dorso-lateral PFC |
| 4 | 28, -47.3, 26.8 | 675 | L. dorso-lateral PFC |
| 5 | .9, 42.9, 50.6 | 675 | L. precuneus |
| 6 | 25.1, 63.1, -3.5 | 648 | L. fusiform gyrus |
| 7 | 1.2, 80.8, 38.9 | 648 | L. precuneus / parieto-occip. fissure |
| 8 | 31.9, -20.1, -12.1 | 567 | L. anterior insula |
| 9 | 6.1, -33.1, 18.4 | 567 | L. anterior cingulate cortex |
| 10 | -1.1, 89, 12 | 540 | R. visual cortex / V1-2 / BA 17-18 |
| 11 | 20.5, 41.3, -11.2 | 486 | L. parahippocampal gyrus |
| 12 | -47, 62.3, 18.2 | 459 | R. temporo-parieto-occip. junction |
| 13 | 14.1, 52.8, 15 | 432 | L. precuneus |
| 14 | 4.3, 30.1, -9.2 | 405 | Midbrain |
| 15 | -32.9, 38.8, -12.7 | 351 | R. parahippocampal gyrus |
| 16 | -4.2, 57.2, -32.7 | 297 | R. cerebellum |
| 17 | 11.6, -48.6, -7.9 | 297 | L. medial PFC |
| 18 | 5.5, 10.8, -10.7 | 243 | L. ventral tegmental area |
| 19 | -1.5, -9.1, -11.4 | 135 | Subcallosal (parolfactory) area / BA 25 |
| 20 | 6.9, -40.9, 35.4 | 135 | L. superior medial frontal gyrus |
